# Improving the Capstone Biochemistry Lab and Evolution to a Course-Based Undergraduate Research Experience: Lessons Learned from the COVID-19 Online Modality

**DOI:** 10.1101/2024.11.11.623056

**Authors:** Alberto A. Rascón

**Affiliations:** San José State University, One Washington Square, San José, CA, 95112, United States; School of Molecular Sciences, Arizona State University, Tempe, Arizona, 85281, United States

## Abstract

The restructuring of an upper division biochemistry lab capstone course intended for biochemistry students with a range of laboratory experience was explored. A goal of the course was to give students practice with necessary skills in biochemical and biological techniques, especially for an entry level general position in biotechnology. The immediate impact of the online capstone course mandated by the COVID-19 pandemic limited students on learning essential hands-on research skills but evolved during the transition back to in-person instruction to include more elements of in-person practice. This article highlights the evolution of the capstone biochemistry lab to an in-person CURE capstone lab, with lessons learned and resources successfully used during the COVID-19 remote course. These include changes in the way information was disseminated, access to online resources, and modifications in student assessments. This article documents how course evolution resulted in a shift in pedagogical strategies leading to building a community of biochemistry learners that could be used to help college faculty in developing a CURE capstone lab.

## INTRODUCTION

San José State University (SJSU) is a large comprehensive primarily undergraduate institution located in the heart of Silicon Valley where it serves a highly diverse student population. SJSU was named the #1 transformative college in the United States by Money magazine in 2020 (1) and ranks #3 in top public schools in the West (2). Many biochemistry students graduating from the Department of Chemistry at SJSU often seek jobs in the Bay Area biotech industry. In the department, there are only two major biochemistry lab courses providing essential techniques needed to successfully secure a position in industry. The first is the introduction Biochemistry Lab (Chem 131A) where students get exposed to pipetting techniques, activity assays, and learn to use UV-VIS spectrophotometers, among other important equipment and techniques related to biochemical experimentation. The second class, taken after Chem 131A and normally towards the end of graduation, is the Biochemistry Lab Capstone (Chem 131B). The capstone lab was developed to train students in an independent research project to help expand on their accumulated undergraduate background knowledge and research skills. These two course offerings provide seniors with a hands-on biochemistry research experience before graduating (typical classes sizes range from 16 to 20 students a semester). However, prior to 2013, Chem 131B was not taught in a consistent manner and student experience was solely dependent on the research interests of faculty assigned to teach the course. Students were not provided with any detailed experimental strategies or goals for the semester, and the information in the syllabus was lacking any detail, with no expectations or learning outcomes. Chem 131A was more structured, offering consistent experiments that touched on the skills needed to prepare students for the independent work in the capstone lab.

Chem 131B course transformation began prior to the pandemic. As a new assistant professor starting in 2013, taking over the class with minimal lab course teaching experience, was daunting. In careful review of the literature on biochemistry laboratories, revealed that many students already do not receive formal laboratory research training as undergrads (3,4). Therefore, as a new professor, it was important to involve students in an authentic research experience, one where they would be allowed to think, work like a scientist, and help develop their scientific identity (5). To do this, the research of *Aedes aegypti* mosquito proteases (6-8), was introduced into the class but in such a way to nurture the undergraduate student research training experience (9-11). Being at a PUI with minimal research infrastructure, faculty must find creative ways to start and maintain a research program that is inviting and nurturing to undergraduate students while helping them grow in their research development (12). Therefore, incorporating a course-based undergraduate research experience (CURE) would not only fulfill the capstone requirements set by the University and department, it would also provide students with an authentic research experience (13) and allow for data collection for grants or manuscripts. Further, as a minority scientist, it is difficult to develop a scientific identity and maintain that identity while taking undergraduate science lecture and lab courses when there is no minority faculty representation (4). By incorporating an “authentic” research experience in Chem 131B, students from diverse backgrounds (academically and socially) would be trained and engaged in a lab setting similar to how professional scientists are trained to help them develop and maintain their own scientific identity (5).

## METHODS

### Initial Development of the Capstone Lab (Chem 131B) Pre-Pandemic

In developing the capstone course, the goal was to allow students to conduct independent research on an important topic that was relevant to not only the population in the area but would have global implications. Therefore, research on understanding the roles of digestive midgut proteolytic enzymes from the *Aedes aegypti* mosquito was used. The mosquito is a known vector of potentially deadly viruses to humans (14), and a direct correlation between protease activity and egg production was found, but the exact role proteases play in this process is still relatively unknown (8,15,16). So, to engage students in an authentic research experience, student projects focused on the cloning and bacterial recombinant expression of these essential enzymes. In addition, the capstone course is a culmination of all biochemistry, molecular biology, and microbiology courses that serve as either pre- or corequisites for the course. Therefore, in structuring the class, it was essential to tie in all the different techniques and background information learned from previous classes into Chem 131B. More importantly, however, it was necessary to focus on developing essential laboratory techniques (*e*.*g*., pipetting, preparing solutions and buffers, lab safety, proper biohazard disposal, and maintaining a proper lab notebook) to then be able to have students learn the molecular biology techniques needed to clone the mosquito protease genes of interest into a bacterial expression vector (**Figure 1**).

**Figure 1.**
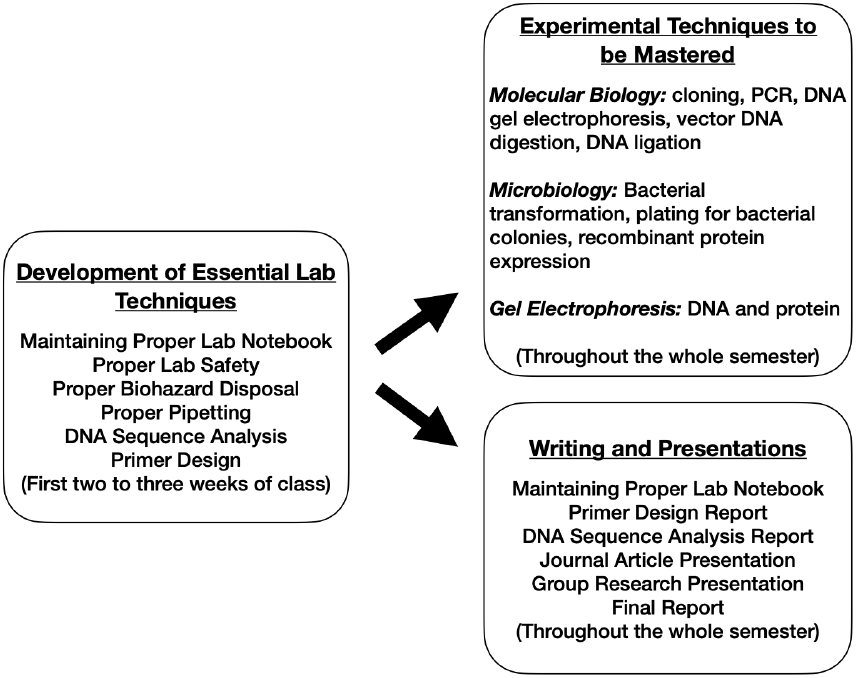
Development of essential lab techniques taught throughout the semester in the Chem 131B Biochemistry Lab Capstone Course. The first two/three weeks are dedicated to building the foundation leading to mastering of experimental techniques, and by the end of the semester, also becoming proficient at writing and presenting scientific work.

Additionally, microbiological techniques were also incorporated to teach students proper aseptic technique, transformation, recombinant protein expression, bacterial culture growths, and other important techniques (*e*.*g*., DNA and protein gel electrophoresis, DNA sequence analysis, to name a few) needed as check points to ensure proper cloning of the genes of interest. Many of the mosquito midgut enzymes used for the course had never been cloned or isolated, giving students an opportunity to be the first in doing so, and as an important incentive, students who were successful and contributed to the projects, would earn authorship on future manuscripts (see the acknowledgement section in Nguyen *et al*. 2018 (8) where four former Chem 131B students earned co-authorship from work conducted in the course). In addition to training students on research techniques, the course was also designed to enhance oral and written science communication. Students were instructed to select a journal article that was relevant to mosquitoes, viruses, or a closely related topic, and with guidance from the instructor (through one-on-one meetings and a rubric), present to the class. The goal was to help students learn to dissect and be critical of published articles, but also help students learn how to properly search for relevant sources that would be needed for their final research paper. As a final component in their training, students were tasked with writing a formal research paper (in the ACS Biochemistry format style) on the work completed throughout the whole semester. A detailed rubric was given outlining important components required for a successful paper (please see Supporting Information), with drafts of each section due throughout the entirety of the semester. Along with the final research paper, students were tasked with presenting the data collected as a mini formal seminar. All of this was used to assess student understanding and progress throughout the semester. Furthermore, different assignments (*e*.*g*., primer design, DNA sequence analysis, drafts of the research paper) were given to be completed outside of class, physical lab notebook carbon copy entries were collected, and a midterm and a final exam were given. The whole approach in training and preparation was in line with the Department of Chemistry Program and Course Learning Outcomes and the University’s General Education requirements (**Table 1**).

**Table 1.** Program, Course, and General Education Learning Outcomes Covered in the Biochemistry Lab Capstone Course (Chem 131B).

| **Program Learning Outcomes (PLOs)** | **Course Learning Outcomes (CLOs)** | **GE Area R (Earth and Environment) Goals** |
| --- | --- | --- |
| **Core Chemistry Ideas (Fundamentals)** |  |  |
| Students will be able to identify, formulate, and solve a range of chemistry problems (fundamental to complex) through application of mathematical, scientific, and chemical principles (PLO 1.1). | Apply proper laboratory practices including safety, waste management, and record keeping (CLO 1). | Students apply knowledge of scientific theories and concepts as well as quantitative reasoning to explore the relationship between humans and the natural environment. |
| Students will be able to recognize, relate, and/or apply chemistry terms and concepts to propose and solve interdisciplinary and multidisciplinary real-world problems (PLO 1.2). | Use and understand modern biochemical techniques and instruments (CLO 2). | Students achieve an understanding of the role that science plays in addressing complex issues, as well as the potential limits of scientific endeavors and the value systems and ethics associated with scientific inquiry. |
| **Experimentation/Lab Practice** | Plan, design, and execute experiments based on biochemical literature (CLO 3). | **GE Learning Outcomes (GELOs)** |
| Students will be able to develop an experiment to address a hypothesis using literature and execute the planned experiment using standard chemistry techniques (PLO 2.1). | Interpret experimental results and draw reasonable conclusions (CLO 4). | Apply scientific principles and the scientific method to answer questions about earth, the environment, and sustainability while recognizing the limits of both the method and principles (GELO 1). |
| Students will be able to acquire, record, and critically evaluate data through use of instrumentation and software, appropriate record keeping practices, figure preparation, and scrutiny of experimental results (PLO 2.2). | Communicate effectively through written and oral reports (CLO 5). | Apply mathematical or quantitative reasoning concepts to the analysis and generation of solutions to issues of earth, the environment, and sustainability (GELO 2). |
| Students will be able to recognize and assess laboratory hazards, practice risk minimization, and conduct safe laboratory practices (PLO 2.3). |  | Communicate a scientific finding, assertion, or theory to a general audience with the integrity and rigor of the underlying science (GELO 3). |
| **Community, Social, Societal Implications** |  | Explain ethical, social, and civic dimensions of scientific inquiry (GELO 4). |
| Students will be able to explore, critique, and reflect on how chemistry relates to society, culture, and issues of equity and ethics that shape their scientific beliefs and identities (PLO 3.1). |  |  |
| Students will be able to identify as scientists within the scientific community through constructing peer reviews, engaging in collaborations, and participating in mentorship (PLO 3.2). |  |  |
| **Communication Skills** |  |  |
| Students will be able to design and deliver engaging presentations on diverse chemistry topics in a professional manner and with clear, concise organization that demonstrates mastery of the topic (PLO 4.1). |  |  |
| Students will be able to integrate research findings into a concise original written report that either analyzes collected data and obtained results or reviews and reflects on published scientific work (PLO 4.2). |  |  |
| Students will be able to identify an audience and construct a message tailored to that audience and act as a science ambassador by conveying the importance of the research or topic of study (PLO 4.3). |  |  |

### COVID-19 Online Modality (During Pandemic)

With the onset of the pandemic, the in-person hands on laboratory-based course had to be altered to a complete online experience, with many asking how this course can be given as an online class, but also, one that meets the Program and Course Learning Outcomes, as well as the General Education Learning Outcomes (GELO) (**Table 1**). The biggest challenge was in maintaining the “authentic” research experience and allowing students to stay engaged and develop their sense of scientific identity. Even within a year of the pandemic and online courses, students were already fatigued with the strictly online modality (17). For the complete online Spring 2021 class, options were explored on how students from Chem 131B, working remotely, could still benefit from working on a midgut protease that no one had isolated or studied *in vitro*. In general (pre-pandemic), the first couple of days in Chem 131B, students were trained and tasked to design primers for the cloning of the mosquito protease gene into a bacterial expression vector, which then, based on their design and guidance, were sent for synthesis. For this approach to work for the online class, it was important to communicate to students that although they could not perform the hands-on experiments in the physical lab, all the work they conducted through video and online, would lead to the cloning and expression of their mosquito protease gene. To connect the online students to experimentation and build awareness of the techniques employed in their work, video recordings of the exact experimental procedures were provided. For example, we developed an awareness and shared fundamental techniques such as pipetting and loading a gel as authentically performed during experimentation. These videos were made available through the Learning Management System (LMS) – CANVAS. In addition, guidance for essential fundamental practices were also made available through video. These included information on how to record research methods in a laboratory notebook, the process of experimentation to presentation, and from experimentation to the final research paper. This was noteworthy because prior to the pandemic none of the material was recorded and no one ever thought about the kinds of resources students would need to build connections between the work they were doing in lab to construct the products that leads to communication of this work. These extra resources proved to be valuable for scaffolding the scientific practices students develop in a lab setting. Furthermore, because of the pandemic, quizzes were implemented as checkpoints to gage student learning and to address misunderstandings. This was important because studies show that students who receive quizzes perform much better than those who do not, showing a correlation between quiz scores and understanding (18). This formative assessment approach helped identify steps to help improve student learning while being remote. Pre-pandemic no quizzes were given, so most student understanding was based on the content of their notebooks, especially since detailed information was asked to be included. For example, if the experiment was PCR, students were tasked with providing background information on the technique (*i*.*e*., the contents needed for successful PCR amplification: primers, DNA template, polymerase), then tie this information to the mosquito gene being amplified and why it is essential to have the proper primers and conditions needed for annealing. Unlike other lab courses, the structure of the notebook in Chem 131B is such that students were required to provide detailed Introduction/Background on the experiment, followed by providing context and tying this to the work being conducted in the lab. They were also required to provide detailed Materials and Methods (with company, catalog numbers of any reagent or biologicals used), Results (with a narrative, figures, and figure captions), a Discussion (with interpretation of the Results obtained), and Conclusion. This approach helped assess the groups that were lacking understanding or needed reinforcement, and although the notebook was utilized as a formative assessment tool, students pre-pandemic did not perform as well as expected on midterms or final exams (**Table 2**). It is noteworthy to indicate that with the video recorded resources available to the online class, and saving the notebook online as a Google Doc (rather than carbon copies), the online students provided more detail and information in their notebooks compared to classes pre-pandemic. Additionally, the online students were engaged in much more meaningful discussions and performed just as well, if not better overall, in oral, writing, and summative assessments (**Table 2**). There were challenges with the online class, but mainly due to ZOOM/remote fatigue (17). All students in class shared that they were tired of being at home, especially with taking all their classes online. Additionally, fellow faculty colleagues shared that many of their students were having similar sentiments about online remote learning. To help overcome this early in the semester, lectures were shortened (from 1.25 h to 45 min). Also, to entice students in keeping their cameras on during lectures, class discussions, and in-class assignments, questions were strategically asked (written prior to the online class meeting) to ensure individual participation. Students could earn anywhere from 1 to 3 points (depending on the difficulty of the question) that counted towards their final grade. Many of the resources that were developed for this online course had not been part of the teaching strategy used prior to the pandemic and the students benefited from having the extra resources to help them learn the material.

**Table 2.**
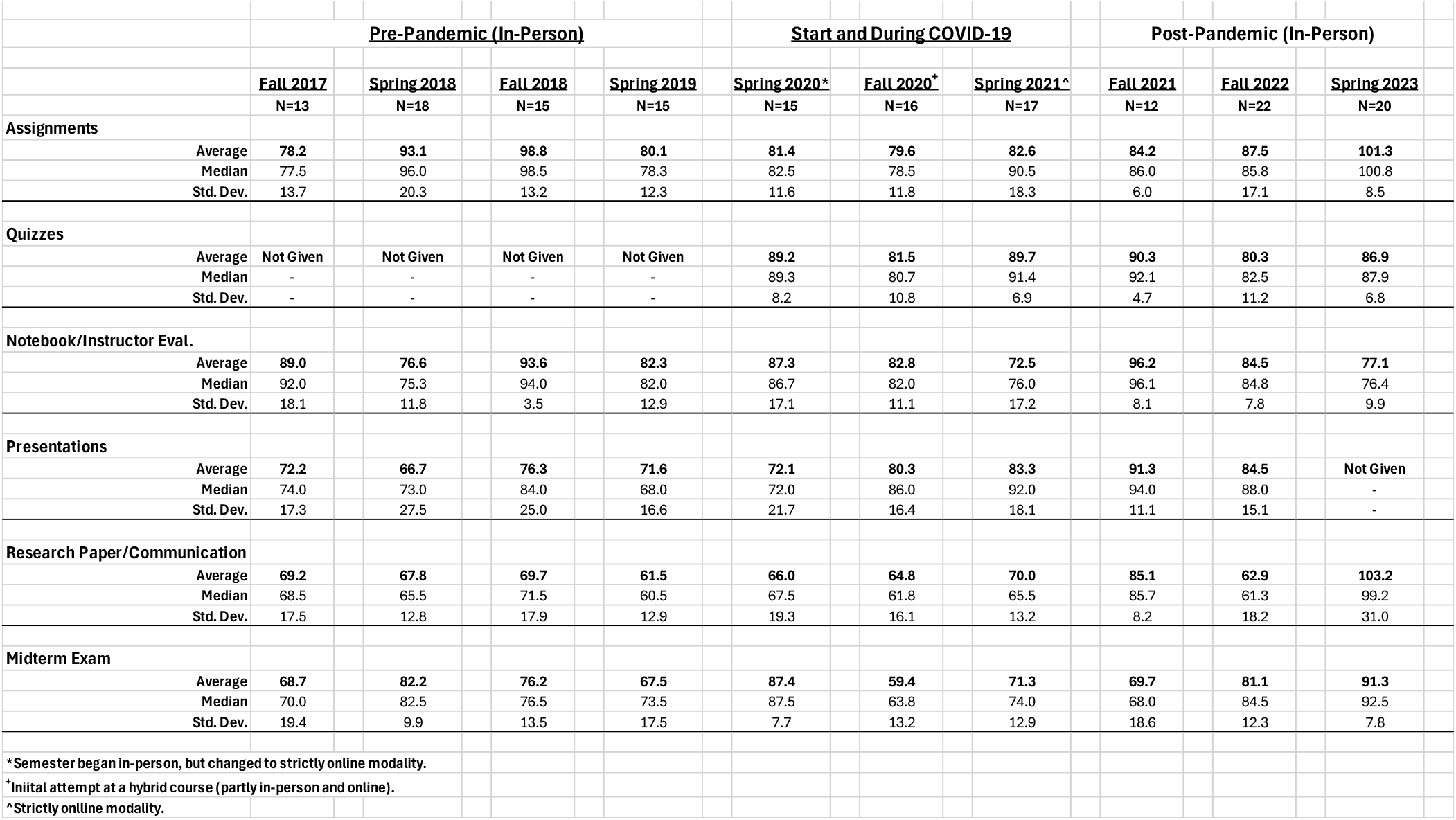
Percentages Earned in Oral, Writing, and Summative Assessments in Chem 131B (before, during, and post-COVID 19 pandemic semesters).

### Evolution of the In-Person CURE with Changes Learned from the COVID-19 Online Modality

Teaching the online lab, led to the realization that the in-person lab could be greatly improved to better help students learn how techniques are used in the context of actual experimentation. This proved to be the most important, if not essential process, since many students (like the online Chem 131B class) were not exposed to or had minimal hands-on lab experience because of the pandemic (19). In fact, many biotech companies were in search of newly graduated students who had hands on research experience but were unsuccessful due to the limitations imposed by the COVID-19 pandemic. Therefore, the approach to improving the in-person class changed keeping in mind the lessons learned from the online Chem 131B class (**Table 3**). The first major change was providing lecture slides and notes, the experimental procedure videos, and other relevant material a week before the actual experiment to activate background knowledge. For example, if PCR was the technique covered, the experimental videos showing how to perform PCR were provided, along with lecture slides and notes covering the material, while emphasizing that although the gene and information modeled in the videos was not of their mosquito protease gene, the experimental setup is nearly identical. This provided context and familiarity to students. In addition, during class enough background information was provided on the technique that students were going to learn, that they themselves would be able to determine what aspects were different between the video and their own experimental setup. As an example, mosquito protease genes contain different percentages of GC and AT content, and depending on the differences (for example, higher GC content than AT) the melting temperature of the designed primer would have to be adjusted, while also needing to adjust the annealing temperature conditions needed to successfully amplify the gene of interest. This video assisted approach served to prepare the class beforehand to allow more time to be spent on conducting hands-on experiments. Students were empowered to try the technique as covered in the video, while providing the flexibility in allowing students to make mistakes and encouraged to reflect on what went wrong and try again. This type of student is known as a Converger in Kolb’s learning cycle (20). Convergers flourish by doing and process information through application, and by providing the “doer” approach in class, these students are more engaged and excited about the work. Other students (known as Assimilators in Kolb’s learning cycle), who rely on lecture slides and notes being available beforehand, as well as any other pertinent material that provides a logical structure (*e*.*g*., experimental videos and approaches) were provided with materials to also be fully engaged and successful in class (20). With the importance of studying and focusing on the human viral vector the *Ae. aegypti* mosquito, provided the Divergers (those students who need to make connections to their own personal lives, interests, or experiences (20)) the motivation to learn and be engaged with the project. With the presence of the *Ae. aegypti* mosquito in the United States, and with the diverse background of many students from areas endemic with the mosquito, made the topic more appealing. Lastly, being the first ever student researchers to work on a never studied or isolated mosquito protease gave the Accommodators (students that learn by applying their problem-solving skills in situations that can lead to self-discovery (20)) a chance to try different approaches, optimize a procedure, or contribute to changes in the experimental setup and improve lab experiments. Given that the structure of the course is designed for students to work in groups of four, in many instances all four types of learning styles would end up in the same group, leading to independence, working cohesively, motivating, and helping each other in the doing (Convergers), feeling (Divergers), reasoning (Assimilators), and discovery (Accommodators) (20).

**Table 3.** Changes in Assigned Material for Chem 131B. Shown are Chem 131B before, during, and after COVID-19 Pandemic

| **<u>Chem 131B Before Pandemic (In-Person)</u>** | **<u>Chem 131B During Pandemic (Online)</u>** | **<u>Chem 131B After Pandemic (In-Person)</u>** |
| --- | --- | --- |
| **Assignments** (Primer design report, DNA sequencing analysis report, report drafts). Meets PLO 1.1-1.2, 2.1-2.3, 3.1-3.2; CLO 1-3; and GELO 1-4). | **Assignments** (Primer design report, DNA sequencing analysis report, report drafts). Meets PLO 1.1-1.2, 2.1-2.3, 3.1-3.2; CLO 1-3; and GELO 1-4). | **Assignments** (Primer design report, DNA sequencing analysis report, report drafts). Meets PLO 1.1-1.2, 2.1-2.3, 3.1-3.2; CLO 1-3; and GELO 1-4). |
| **Lab Notebook (physical carbon copies) and Instructor Evaluation.** Meets PLO 2.1-2.3; CLO 1-4; and GE Area R writing requirement. | **Lab Notebook (online Google Doc) and Instructor Evaluation.** Meets PLO 2.1-2.3; CLO 1-4; and GE Area R writing requirement. | **Lab Notebook (online Google Doc) and Instructor Evaluation.** Meets PLO 2.1-2.3; CLO 1-4; and GE Area R writing requirement. |
| **In-class Midterm Exam.** Meets PLO 1.1-1.2, 2.1-2.3, 3.1-3.2; CLO 2, 4; and GELO 1-4. | **Online Midterm Exam.** Meets PLO 1.1-1.2, 2.1-2.3, 3.1-3.2; CLO 2, 4; and GELO 1-4. | **Take-home Midterm Exam.** Meets PLO 1.1-1.2, 2.1-2.3, 3.1-3.2; CLO 2, 4; and GELO 1-4. |
| **Journal Article and Group Research Presentations.** Meets PLO 4.1, 4.3; GELO 3-4. | **Journal Article and Group Research Presentations.** Meets PLO 4.1, 4.3; GELO 3-4. | **Research Communication (ACS Biochemistry style).** Meets PLO 3.1, 4.2-4.3; CLO 1-4; and GE Area R writing requirement |
| **Full Research Paper (ACS Biochemistry style).** Meets PLO 3.1, 4.2-4.3; CLO 1-4; and GE Area R writing requirement | **Research Communication (ACS Biochemistry style).** Meets PLO 3.1, 4.2-4.3; CLO 1-4; and GE Area R writing requirement | **Online Quizzes (8 total, lowest scored dropped).** Meets PLO 1.1-1.2, 2.1-2.3, 3.1-3.2; CLO 1-4. |
| **In-class Final Exam.** Meets PLO 1.1-1.2, 2.1-2.3, 3.1-3.2; CLO 1-3; and GELO 1-4). | **Online Quizzes (8 total, lowest scored dropped).** Meets PLO 1.1-1.2, 2.1-2.3, 3.1-3.2; CLO 1-4. |  |

## RESULTS and DISCUSSION

Prior to the pandemic, limited resources were made available to students in Chem 131B, lecture times were as long as 1.25 h, and the only formative assessment tool used was the midterm exam. During the pandemic, students were not able to conduct hands-on research, so lecture times were reduced to 45 min, and online resources (*e*.*g*., pre-recorded experimental procedures and lecture videos), as well as quizzes were incorporated. Post-pandemic, students missed out on gaining the hands-on research skills needed to be successful in in-person lab, but also to be successful in securing a position in biotech industry. Therefore, lectures were condensed further to 30-40 min and were provided even more online resources (*e*.*g*., live recorded lecture videos, recorded experimental procedures, results, class discussions) to focus more on in lab hands-on research techniques. In addition, quizzes and online notebooks were used as formative assessment tools to gauge student understanding. These changes to the authentic CUREs approach not only allowed students to develop and maintain their scientific identity they also improved their overall understanding of the material leading to percent increases in three major areas: Assignments, the Midterm Exam, and Research Paper/Communication (**Table 2**). Students post-pandemic took ownership of the project, immersed themselves in different approaches when cloning was not working or when soluble expression was not achieved because there was more in class time to do so. Due to troubleshooting of experiments and delving more into the literature to find alternatives to an experimental approach, increased their overall understanding. The notebook entries were more detailed than pre-pandemic and during pandemic classes, their confidence in having meaningful discussions with the instructor and each other increased, and their quiz/midterm exams were some of the highest earned (**Table 2**), which led to better oral and written communication. The new approach and the tools provided for success were brought by the new revamped curriculum. Although it is difficult to compare post-, pre-, and during pandemic students with each other, given that different pedagogical approaches were taken, the evolution of Chem 131B post-pandemic led to positive changes in student outcomes. With more resources, along with measurable formative assessment tools, provided in the moment data needed to help modify, reenforce, or review concepts not understood during class. This led to substantial improvement and understanding as shown in the summative assessment (the final research/communication paper) (**Table 2**). In classes prior to and during the pandemic, assessment tools were not as efficiently used, which possibly led to the lower overall percentages in the final summative assessment and final grades.

Learning from Kolb’s experimental learning cycle (13,20), led to the realization that students from the online course (during the pandemic) understood the mosquito work better than pre-pandemic students because of the transformative approach taken to deliver the material and expose students to a real-time research experience, even though they were unable to conduct the experiments themselves. The pandemic forced changes in providing recorded material and extra resources, as well as the use of different assessment tools, and provide inventive ways to keep the class engaged. In returning to in-person lab after the mandate was lifted, the entire curriculum was restructured to help train students more efficiently and give them a better hands-on experience, by focusing more on their lab techniques, their written and oral communication, and shortening lecture times. Further, revamping the in-person CURE capstone lab and curriculum resulted in an unexpected outcome, a community was built that provided a safe and comfortable learning environment. Students have often commented on how scared and intimidated they were to ask questions at the start of the semester, especially given that many do not have a formal research experience and felt inadequate to engage in scientific conversation. The fact was that students were unaware of the new approach of how Chem 131B was going to be taught and how they were going to be led into an independent research experience, so they would often acknowledge that the course was going to be difficult. Once students realized that the goal was to allow them the opportunity explore different techniques, optimize different approaches, but more importantly, allow them to make mistakes and be comfortable with failing, did more students buy into the learning approach and the independent project. In Figure 2, are paraphrased student comments from Spring 2021, Fall 2022, and Spring 2023, highlighting the feelings of the students in class. This was very important because the pandemic revealed many inequities and struggles of many college students, which lead to hinderances in student learning and potentially delaying graduation (21,22), but by allowing open communication, and importantly, also sharing career struggles as a minority first generation student from a low-income socioeconomic background, students felt more comfortable in sharing their overall college and post-pandemic struggles. The goal of revamping Chem 131B was to provide the best CUREs for students not getting the hands-on experience needed before graduating (exacerbated by the pandemic), and in doing so, the unintended consequence was a safe learning and collaborative environment that allowed students to share their struggles with each other and the instructor. A major challenge faced by many students during the transition from in-person to online learning was finding a suitable substitute for the social aspect of learning and working together (23). By making the course more engaging and leaning towards more collaborative work, students felt much more comfortable and confident in their scientific abilities. In addition, many students in Chem 131B are first-generation from low to middle socioeconomic backgrounds who normally come from nonacademic backgrounds and may not be familiar with the research environment, leading to feelings of isolation or intimidation of research and research advisors (24,25). This isolation or fear of science keeps many students from pursuing higher degrees or prevents students from approaching research opportunities. Therefore, by taking this approach in revamping the CURE course, helped students overcome their insecurities, allowed growth and positive, more student-centric changes to the curriculum, and in the future may lead to diversifying science (24,25).

**Figure 2.**
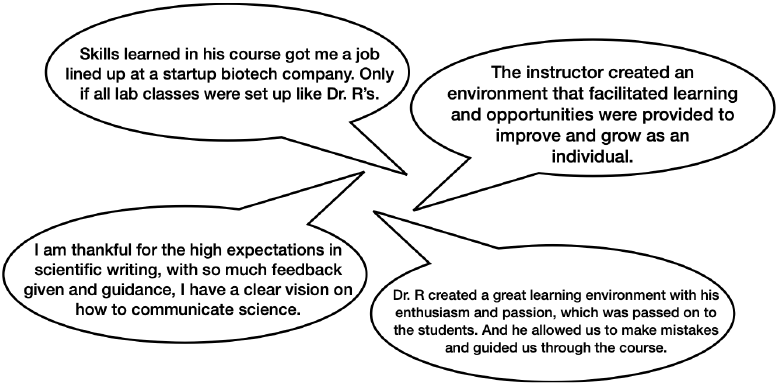
Paraphrased overall comments from Chem 131B students in Spring 2021, Fall 2022, and Spring 2023.

## CONCLUSIONS

The COVID-19 pandemic allowed technology to shine, allowing educators to create new ways to incorporate high-impact learning online and adequate communication (18,19,26-30). However, many students lost the opportunity to gain hands-on lab and research training, leading to difficulty in learning proper lab techniques. Here, the Chem 131B capstone lab course post-pandemic curriculum was revamped to provide students more in class time to not only help improve their overall hands-on lab experience but their overall understanding of the mosquito protease project. The CUREs approach helped students develop essential research skills in biochemistry. Additionally, providing recorded lecture and experimental procedures, as well as providing extra resources for better understanding of the material, led to students taking ownership of the in-class project, helping students develop and maintain their scientific identity, while giving them confidence in oral and written science communication. An unexpected consequence of the new revamped class was the creation of a community, where students felt safe and comfortable with failing. Students worked at their own pace, modifying, or optimizing steps that helped in cloning a mosquito protease gene into a bacterial expression vector. Taken together, learning from the COVID-19 online experience helped improve student learning and training post-pandemic, making the course-based undergraduate research experience more fruitful and engaging. This work could be used to help college faculty in developing a more hands-on CURE capstone lab. In the future, formal student assessment studies are needed to further strengthen the ideas highlighted in this work.

## Supporting Information

Chem 131B Lab Research Communication Grading Rubric (PDF)

## ACKNOWLEDGMENTS

I would like to thank Dr. Resa Kelly (Dept. of Chemistry and Science Education, San José State University), Dr. Anita Nag (Division of Natural Sciences and Engineering, University of South Carolina Upstate), and Ms. Flor S. Cisneros (Substitute Teacher and California State Teach Credential Graduate) for their helpful feedback and suggestions. The *Aedes aegypti* mosquito work incorporated into the capstone course was supported by the National Institute of General Medical Sciences (NIGMS) of the National Institutes of Health (NIH) under Award Number SC3GM116681 to AAR while at San José State University. Special thanks go to all my former Chem 131B students, who without them, I would not have evolved to become a better teacher.

