## Supporting Info for "Improving the Capstone Biochemistry Lab and Evolution to a Course-Based Undergraduate Research Experience: Lessons Learned from the COVID-19 Online Modality"

Alberto A. Rascón, Jr.*

School of Molecular Sciences, Arizona State University, Tempe, Arizona, 85281, United States

**<u>Chem 131B Lab Research Communication Grading Rubric</u>**

Points will be assigned to the abstract, figures/figure captions, and the body of the Communication to give the final combined total score. Each section is weighted differently, with the most points awarded to the ***Body of the Communication (40 pts)***, followed by the ***Abstract (10 pts), and Figures, Figure Legends or Tables (10 pts)***. Points will also be given for proper ***Information Literacy & References (5 pts)***, an appropriate ***Title (5 pts)***, and ***Overall Impression*** of the communication ***(5 pts)*** for a ***<u>total</u>*** of ***<u>75 points</u>***.

***<u>Important note:</u> <u>No Materials and Methods section should be included, this should be in the Notebook!</u>***

***<u>The Body of the Final Communication must be 1,000 words in length (not including figure legends, tables, references, etc…). The Abstract is only 250 words. Overall points will be deducted from each section if too long or too short! See info below for each section.</u>***

***The content of the research paper is based and modified from the Biochemistry Communication journal article Author guidelines (ACS Publications) found at: <u>https://publish.acs.org/publish/author_guidelines?coden=bichaw#manuscript_types</u>, and introduced in the class, and will be evaluated as described below:***

***Title (5 pts)*** - The title of the research paper should be brief, informative and self-explanatory. The reader should be able to read the title and be able to surmise the contents of the paper without actually reading the entire paper. The title should not be so technical that only specialists will understand, but appropriate for a general audience. ***Include a title with your Research Communication draft!***

***Abstract (10 pts)*** - The abstract should be equal or less than 250 words, and should succinctly present the work studied, the significance of the work, the experimental approach used, the major findings, and conclusions. It is imperative that the abstract is self-explanatory and suitable to help set up the communication.

***Body of Communication (40 pts)*** - ***This section will be graded on content, clarity, and coherence.*** This section should include the purpose of the investigation and how it relates to other work in the field, providing enough background information to help state why there is interest in this particular field of study, followed by the results obtained. However, the results are presented concisely and with the flow of the text, summarizing the main findings of the work. Along with the results presented, a discussion of experimental results or speculation of why a result was obtained should be included. ***It is important to note that not all data collected throughout the semester should be included.*** Once the figures/tables are presented, use this information to interpret the results ***<u>AND</u>*** to relate them to existing knowledge in the field. Highlight the ***<u>MOST</u>*** significant results. Finish the section with a conclusion/future directions emphasizing why the work is relevant. ***Find enough references to help write this section.***

***Overall, general questions that should be considered to help write this section are:*** Why are we studying the protease? What makes this protease different? Is it the no leader? EK version? Which vector was the gene cloned into, pET28a, pET28b, or pET29b? How do the results obtained relate to the original problem or question? Do the data support the hypothesis given in the introduction? Are the results consistent with observations made by other investigators? If the results obtained are unexpected, explain why, especially using the literature to find appropriate references to help support your statements, assumptions or speculations. What further research would be necessary to answer the questions raised by your results. How do your results fit into the big picture? In addition, propose what the next major experiments are. For example, growth experiments to determine if the protease can be expressed in bacterial cells, then what's next? ***The Body of the Communication should be 1,000 words. So be concise! Don't write any unnecessary information or "fluff"!***

- ***35 - 40 pts*** - The points in this section are ***clearly*** defined and concisely presented. Appropriate background information is given, along with the purpose and interest in this subject. The data collected is cited and its relationship to the conclusion is clear. Most of the questions in the above description are covered and answered. Relevant references (and enough references) are used to help support statements, assumptions, or speculations made. The writing is clear, sentences are complete, words are properly spelled, and more importantly, sentences are well-paraphrased and cited to avoid plagiarism.

- ***29 - 34 pts*** - The points in this section are ***reasonably*** defined and well presented. Appropriate background information is not fully given, the purpose and interest in this subject are not clearly explained. The data collected is not always cited or effectively used to support the conclusion. Not many of the questions in the above description are covered and answered. Relevant references, but not enough references are used to help support statements, assumptions, or speculations made. The writing is legible, but there are ***minor*** problems with grammar, spelling, and/or punctuation. Minor issues with paraphrasing and properly citing sentences, but no obvious signs of plagiarism.

- ***23 - 28 pts*** - The points in this section are ***not*** well presented. Although most points are present, some are missing. Terms and ideas are given, but are not discussed or defined. Appropriate background information is not given, the purpose and interest in this subject are not explained. The data collected is not cited or used to support the conclusion. Questions in the above description are not well covered or answered. Even though statements, assumptions or speculations are made, there is a lack of relevant references to support them. The writing is legible, but there are obvious problems with grammar, spelling, and/or punctuation. There are issues with paraphrasing and properly citing sentences, but no obvious signs of plagiarism.

- ***17 - 22 pts*** - The points presented in this section are not effectively made. Appropriate background information, purpose, and interest in this subject are not given. Terms and ideas are hardly given, discussed or defined. Questions in the above description are hardly covered or answered. Major lack of relevant references, even though statements, assumptions or speculations are made. The writing is barely legible and there are major problems with grammar, spelling, and/or punctuation. There are major issues with paraphrasing and properly citing sentences, but no obvious signs of plagiarism.

- ***11 - 16 pts*** - The points presented in this section are missing. There is lack of background information, purpose, and interest in this subject. Terms and ideas are not given, discussed or defined. Questions in the above description are not covered or answered. Major lack of relevant references, even though statements, assumptions or speculations are made. The writing is barely legible with major problems with grammar, spelling, and/or punctuation. There are major issues with paraphrasing and properly citing sentences, with obvious signs of plagiarism.

- ***0 - 10 pts*** - The information presented is illegible, to the point that discerning what is being written is impossible and difficult to score. No references and/or improper citations are obvious. Improper paraphrasing and plagiarism is obvious.

***Figures, Figure Legends and Tables (10 pts)*** - Figures and tables should be used only to document experimental results or methods that cannot be adequately described in the text. The table and figure legends should include the figure number and a brief description, preferably two to three sentences ***OR*** descriptive phrases. The legends should be understandable ***WITHOUT*** reference to the text and should not include new material that is not found within the text. Explain all symbols and abbreviations used in the illustrations.

***Information Literacy and References (5 pts)*** - This section was given separate points to ensure that proper literature searches were conducted. References are important because they clearly identify the original contributor to the work being cited. The communication requires relevant references to give background information and to introduce knowledge or lack of knowledge in the field of interest. Similarly, the discussion within the communication requires references that will help with interpretation of results, whether they have been observed by other scientists, or if the results are unexpected, appropriate literature searches can be used to explain why this result was obtained.

- ***4.5 - 5 pts*** - Relevant references (and enough references) are used in the introduction and discussion section, and sentences are well-paraphrased and cited to avoid plagiarism.

- ***3.5 - 4 pts*** - Relevant references, but not enough references are used in the introduction and discussion section, and there are minor issues with paraphrasing and properly citing sentences, but no obvious signs of plagiarism.

- ***2.5 - 3 pts*** - There is a lack of relevant references in both the introduction and discussion section, and there are issues with paraphrasing and properly citing sentences, but no obvious signs of plagiarism.

- ***1.5 - 2 pts*** - Major lack of relevant references in both the introduction and discussion section, and there are major issues with paraphrasing and properly citing sentences, but no obvious signs of plagiarism.

- ***0 - 1 pts*** - No references or improper citations are obvious. Improper paraphrasing and plagiarism is obvious.

***Overall Impression (5 pts)*** - These points are given to provide an overall impression of the research paper, taking each section into consideration.
